# Atomic modeling of radiation damage in cryoelectron microscopy datasets

**DOI:** 10.64898/2026.08.21.746204

**Authors:** Alexander Shtyrov, Hugh Wilson, Garib N. Murshudov

## Abstract

Damage to biological specimens by the electron beam is the fundamental resolution-limiting factor in cryoelectron microscopy (cryo-EM) single particle analysis. There is, however, currently no method to accurately infer fluence-dependent changes to the specimen structure during electron irradiation. We develop a Bayesian framework to fit a sequence of atomic models to a series of cryo-EM reconstructions produced at increasing fluence. In particular, our algorithm is able to infer the ensemble average position and atomic displacement parameter of every atom in the macromolecule as a function of fluence. Application of the algorithm to cryo-EM datasets shows that the molecule expands during imaging and identifies environment-dependent variations in beam-induced damage. We use our results to propose a stochastic process model of this phenomenon. We envisage that our method will lead to a better mechanistic understanding of radiation damage to biological specimens and may contribute to efforts to mitigate its effects.

---

The majority of experimentally-determined macromolecular structures are solved by scattering a high-energy beam of X-rays or electrons off a biological sample [7, 41]. Part of the beam scatters elastically, that is, without deposition of energy in the sample — this is the signal that is generally used for structure solution. The remaining part of the beam scatters inelastically, transferring energy into the sample, thus inducing changes such as excitations, ionization and radiation-induced chemical reactions [30]. These changes, whether caused directly by the incident radiation or by secondary reactions, are collectively referred to as radiation damage (RD). RD is a major problem for macromolecular structure determination because it alters or destroys molecules in the sample in a random fashion, thereby reducing the effectiveness of structure determination methods that rely on sample uniformity. In some cases, RD also induces specific chemical reactions in the sample, leading to observation of a state of the macromolecule which does not reflect its biological function.

The present work focuses on cryoelectron microscopy single particle analysis (cryo-EM SPA). Cryo-EM SPA is a technique for reconstructing the electrostatic potential (ESP) of a macromolecule in aqueous solution from projections of this quantity. The projections are obtained from a sample of the macromolecule embedded in vitreous ice imaged in a transmission electron microscope (TEM), where each particle in the sample has a random orientation [38]. Thanks to significant advances in TEM hardware and data processing [31], cryo-EM SPA has joined X-ray diffraction (XRD) as one of the two most widely applied techniques for structure solution. RD is particularly relevant in cryo-EM SPA as it fundamentally limits the total dose that can be used in a single exposure, and hence the signal-to-noise ratio (SNR) of the resulting micrograph. The SNR in cryo-EM SPA is low, placing a lower bound of around 50 kDa on the size of macromolecules that can be studied by this method [57].

Most knowledge about mechanisms of RD to biological samples at cryogenic temperatures comes from XRD studies. RD in XRD may be categorized as either global (affecting the entire crystal) or site-specific (caused by radiochemistry of particular functional groups). Site-specific damage was first reported around the year 2000 [10, 43, 55]. Three radiation-induced reactions appear to be relevant to all proteins at both cryogenic and non-cryogenic temperatures: breakage of disulfide bonds, decarboxylation of acidic side chains, and cleavage of Met to liberate methane or methanethiol [12, 21]. Moreover, the reactions consistently occur in the order listed. Of particular relevance to this study is decarboxylation of Asp and Glu, which has been studied experimentally in some detail. The reaction has the two-step mechanism (1) R– CH_2_COO^−^ **→** R– CH_2_COO^•^ + e^−^, (2) R– CH_2_COO^•^ → R– CH_2_^•^ + CO_2_ [9, 46]. Hallmarks of global RD in XRD include decreased intensity of reflections, as well as increased mosaicity and unit cell volume [20]. A number of empirical models of intensity decay with dose have been proposed (reviewed in [14]). Perhaps the most popular model assumes that the *B*-factors of atoms in the molecule increase linearly with dose [29, 32, 51], implying an exponential decay in the intensity of reflections. Procedures have been proposed which correct the intensity decay during data processing [16].

We now review prior work on RD in cryo-EM SPA and electron diffraction (ED), focusing on biological samples at cryogenic temperatures. The absence of crystal effects makes cryo-EM SPA in particular an excellent model system for studying RD of biomolecules, since we can directly observe beam-induced changes to the molecule without having to account for changes to the structure of the crystal. Several studies report site-specific RD in cryo-EM SPA [5, 28]. Site-specific RD in ED appears to be similar to that observed in XRD [26]. The primary manifestation of global RD in ED is, as in XRD, a fading of reflections which can be modeled by an exponential function of fluence (number of electrons per unit area of the sample) [2, 52]. Experimental observations of the temperature dependence of the fading [3, 49], supported by calculations from Henderson [27], eventually led to the widespread adoption of liquid nitrogen cooling in biological EM. The fluence-dependent decrease in SNR has also been measured using SPA [23]. The *RELION* package for SPA data processing has introduced a weighting scheme to account for this decrease [60], based on fitting a *B*-factor independently to each frame of movies in the dataset. However, *RELION* does not include a model for the change in *B*-factor with fluence.

In this work, we approach RD modeling as an atomic model refinement problem. Briefly, atomic model refinement is the problem of inferring the parameters of a model that best describes the ESP of a molecule as reconstructed from an electron microscopy dataset. The parameters in macromolecular structure determination typically consist of atomic coordinates and *B*-factors, the latter describing the uncertainty in atomic position due to thermal motion and local disorder (see [35] for a review of atomic model refinement). We develop a refinement algorithm for inferring the atomic coordinates and *B*-factors of multiple atomic models from a time series of cryo-EM maps reconstructed at increasing fluence, effectively producing a ‘movie’ of the macromolecule as it is irradiated. We develop our algorithm within a Bayesian framework. The approach allows us to distinguish site-specific RD from noise and the progressive loss of resolution due to global RD. We then apply the algorithm to a series of cryo-EM datasets and propose a stochastic process model of RD. Recent developments in EM techniques, in particular sample supports that minimize beam-induced motion [37], now permit the detailed investigation of RD in cryo-EM SPA by decreasing the effect of other confounding factors on data quality.

## Results

### Proteins expand on exposure to the electron beam

We have developed an atomic model refinement scheme that, when given a time series of cryo-EM maps reconstructed at increasing fluence, infers the parameters of an atomic model describing the macromolecular structure at each time point. To demonstrate its utility, we have applied the algorithm to five high-resolution cryo-EM datasets: catalase enzymes from three species, which we refer to as catalase L, catalase T and catalase H, mouse apoferritin, and *E. coli* DNA-binding protein from starved cells (Dps). Full details of the datasets are given in the Methods. The datasets were collected on a support that minimizes sample motion during imaging, allowing us to focus exclusively on modeling changes to the molecule itself. Details of the datasets are given in the Methods. For each dataset, we obtained a time series of atomic models, each one describing the ensemble average atomic coordinates and *B*-factors after exposure of the sample to a given electron fluence. Together, these models form a ‘movie’ describing the dynamics of the protein as it is progressively damaged by the electron beam. The time series we obtained are a rich source of information about the global and site-specific damage events experienced by the protein, as well as its structure at low dose, before the onset of RD.

We first assessed the effect of the electron beam on the global structure of the protein. In all five datasets, we found that the coordinates of atoms slowly move apart, as shown by the root mean square distance (RMSD) of each atomic model from the first atomic model in the time series (Figure 1A). The RMSD increase is approximately linear at the start of the exposure, and the value of the RMSD appears very similar across the five proteins we examined. We also calculated the RMSD between each model in the time series and the atomic model that we used as a starting model for the algorithm. The latter was refined against the consensus cryo-EM reconstruction (without dose fractionation) using *Servalcat*. In all cases, we found that the first model in the time series was not the one with the smallest RMSD from the input structure, which was instead most similar to a model at a later stage in the time series. However, the RMSD from the first model was small in all cases (with a maximum value of 0.1 Å across the five datasets).

**Figure 1:**
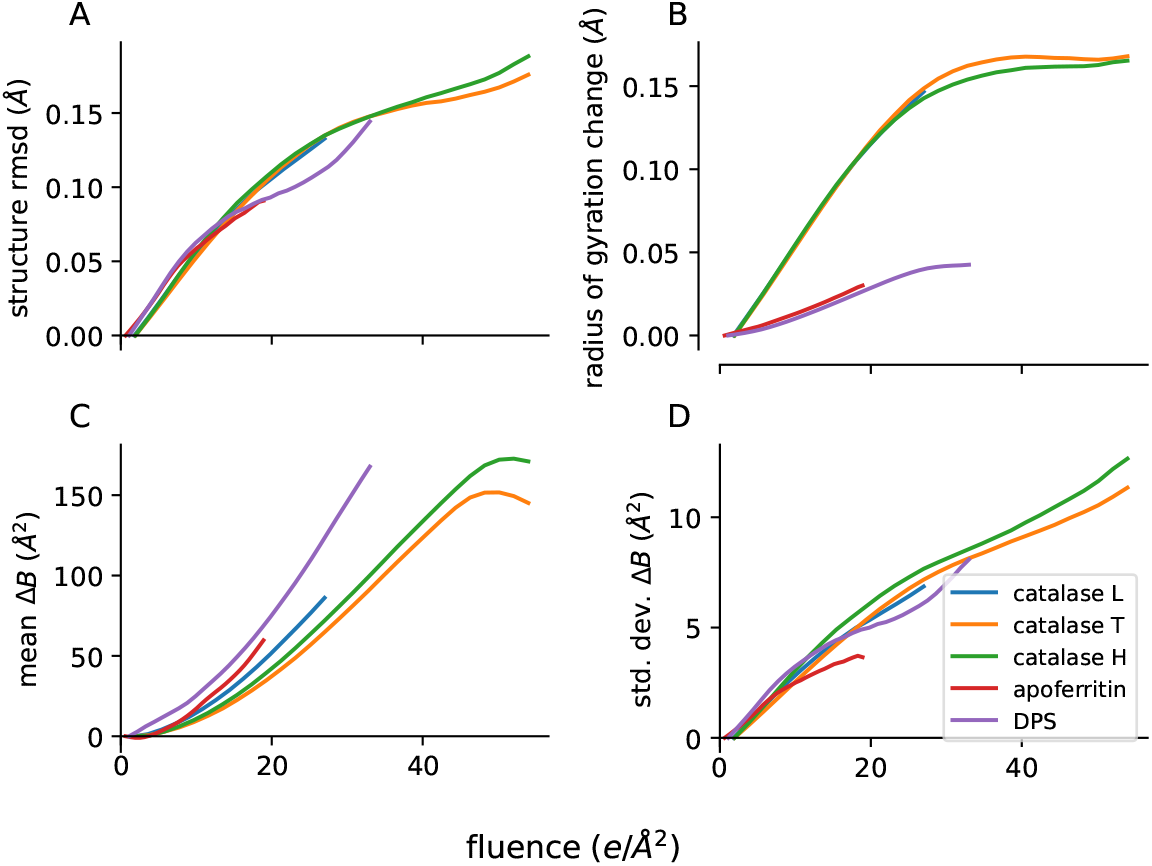
Global metrics of protein behavior under the electron beam. Colored lines indicate different datasets, see legend. (A) Root mean squared distance (RMSD) of atomic model from first atomic model in the time series. (B) Radius of gyration. (C) Mean difference of atomic *B*-factors from their values at the start of the time series. (D) Standard deviation of the difference of atomic *B*-factors from their values at the start of the time series.

While examining the atomic models produced by our program, we observed an apparent expansion of each macromolecule around its center. We quantified this observation by calculating the radius of gyration (RoG), which is the RMSD of the atoms in the protein from its center of mass (Figure 1B). We observed a systematic increase in the RoG as fluence is increased, showing that the molecule indeed undergoes beam-induced expansion. While the absolute difference in RoG is small, we note that it is a very smooth function, and the fact that it is observable in a reconstruction of the ESP averaged over many particles strongly implies that the observed trend reflects the true behavior of the protein. We further validated our results by calculating the RoG directly from the raw cryo-EM maps (Extended Data Figure 1). The maps were first rescaled to have the same power spectrum as the final map in each time series and the absolute value of map values was taken. The trend in the map RoG replicates that in the RoG calculated from atomic models.

### Atomic *B*-factors show marked non-linearity as a function of fluence

Following our analysis of the positions of atoms in the atomic model time series, we turned our attention to the atomic *B*-factors, which describe how disordered the atom is in the structure. Specifically, if the position of an atom is assumed to follow a Gaussian distribution, the *B*-factor is proportional to the variance of this distribution [24]. In this work, we use isotropic *B*-factors, that is, we assume that the Gaussian has radial symmetry about the atom center. RD increases the disorder in a protein by inducing chemical changes in the residues that it comprises [20], leading to heterogeneity among the particles present in the sample. The *B*-factor is therefore an important measure of the extent of RD to atoms in the protein.

First, we calculated the mean and standard deviation of the distribution of *B*-factors at each time point (Figure 1C/D). To ensure that our calculations reflected time-dependent changes in *B*-factors rather than natural variation across the structure, at each time point we used the difference between the atomic *B*- factors and their values in the first atomic model in the time series, which we denote Δ*B*. As expected, the *B*-factors increase with increasing fluence. The standard deviation of Δ*B* also increases. However, we note that both quantities show a marked non-linearity. In particular, the rate of change of the mean appears to increase with time. We later attempt to rationalize this behavior within the framework of a stochastic process model. In order to ensure that the non-linearity is not an artifact of atomic model refinement, we calculated *B*-factors directly from the raw cryo-EM maps (Extended Data Figure 2). We used the formulas in [45] to fit a *B*-factor to each map, relative to the first map in the time series. We compared the obtained values with a straight line fitted to the data by ordinary least squares, and in all cases observed a departure from linearity, replicating the trend observed in the *B*-factors of the refined atomic models.

### Susceptibility of side chains to damage depends on their environment

Next, we focused on RD to specific sites in the protein. Even though RD to proteins is a stochastic process, some radiolysis reactions are known to be particularly favored. Out of the three principal RD reactions, we are only able to investigate decarboxylation of Glu/Asp and cleavage of the C– S bond in Met, since there are no disulfides present in the five protein structures we examine. As in the previous section, we used *B*-factors as a measure of RD to atoms. To better visualize changes in *B*-factors, we removed the global trends identified in the previous section by converting *B*-factors to *z* scores, which we do by subtracting the mean and dividing by the standard deviation at each time point. Figure 2 shows the mean *z* score of atoms in side chains of Asp, Glu and Met. They are shown alongside atom types which are not reported to be susceptible to site-specific damage. Only atoms with a *B*-factor less than the structure median in the first model in the time series were considered in the analysis. This criterion selects for atoms that are ordered at the start of imaging, ensuring that observed trends are due to beam-induced changes rather than side chain disorder already present in the molecule before irradiation. We see that the relative susceptibility of different side chains to damage in cryo-EM SPA is similar to that observed in XRD or ED. Asp/Glu carboxylates are markedly more susceptible than Asn/Gln, confirming the presence of beam-induced decarboxylation. The Met methyl carbon also shows a rapid increase in disorder during electron irradiation, in contrast to Met C*α*. An exception to this trend is the Dps structure, where the two Met C*ϵ* used to calculate the average *z* score appear to remain ordered up to high fluence. Finally, Phe C4 (the carbon atom in the *para* position with respect to C*β*) has a mean *z* less than zero throughout the time series, suggesting this amino acid is resistant to damage. The observed behavior of Phe is consistent with the known RD resistance of organics with conjugated electron systems [44].

**Figure 2:**
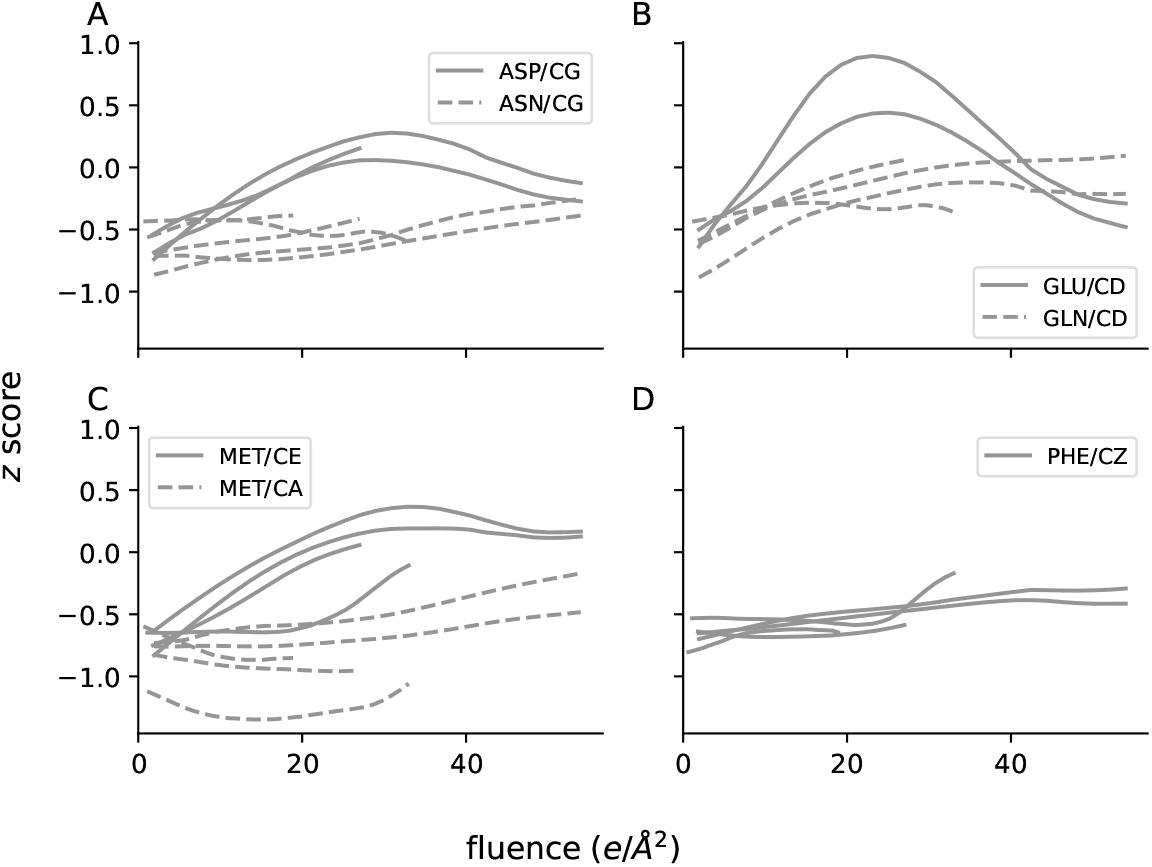
Sensitivity of different amino acids to RD. Each panel shows the mean *z*-score of *B*-factors of atoms of the indicated types. Each line represents the value for a single dataset. Only atoms which have *B* less than the structure median at the start of imaging were included in the calculation. (A) Asp and Asn C*γ*, (B) Glu and Gln C*δ*, (C) Met C*ϵ* and C*α*, (D) Phe C4.

Although we note clear trends in the average behavior of amino acids under irradiation, there is large variation between different residues of the same amino acid present in a protein. We decided to investigate the variability in the extent of damage experienced by Asp C*γ* atoms, since this is a known hallmark of damage in EM and XRD [21]. The catalase structures we analyze in this work are homologs with a high degree of sequence identity, including many conserved aspartates (further comparison of these structures will appear in [48]). The similarity of the three structures allowed us to perform a comparative analysis of the effect of small perturbations in the environment of conserved residues on their susceptibility to damage. Considering the set of *z* scores (one at each time point) for each atom as a vector, we used hierarchical clustering to place all C*γ* atoms in a structure in order from most susceptible to least susceptible to RD. The ordering is visualized as series of heatmaps in Figure 3A–C. There is very significant variation in the behavior of different aspartates, ranging from side chains that are already disordered at the start of imaging to those that are almost entirely resistant to damage.

**Figure 3:**
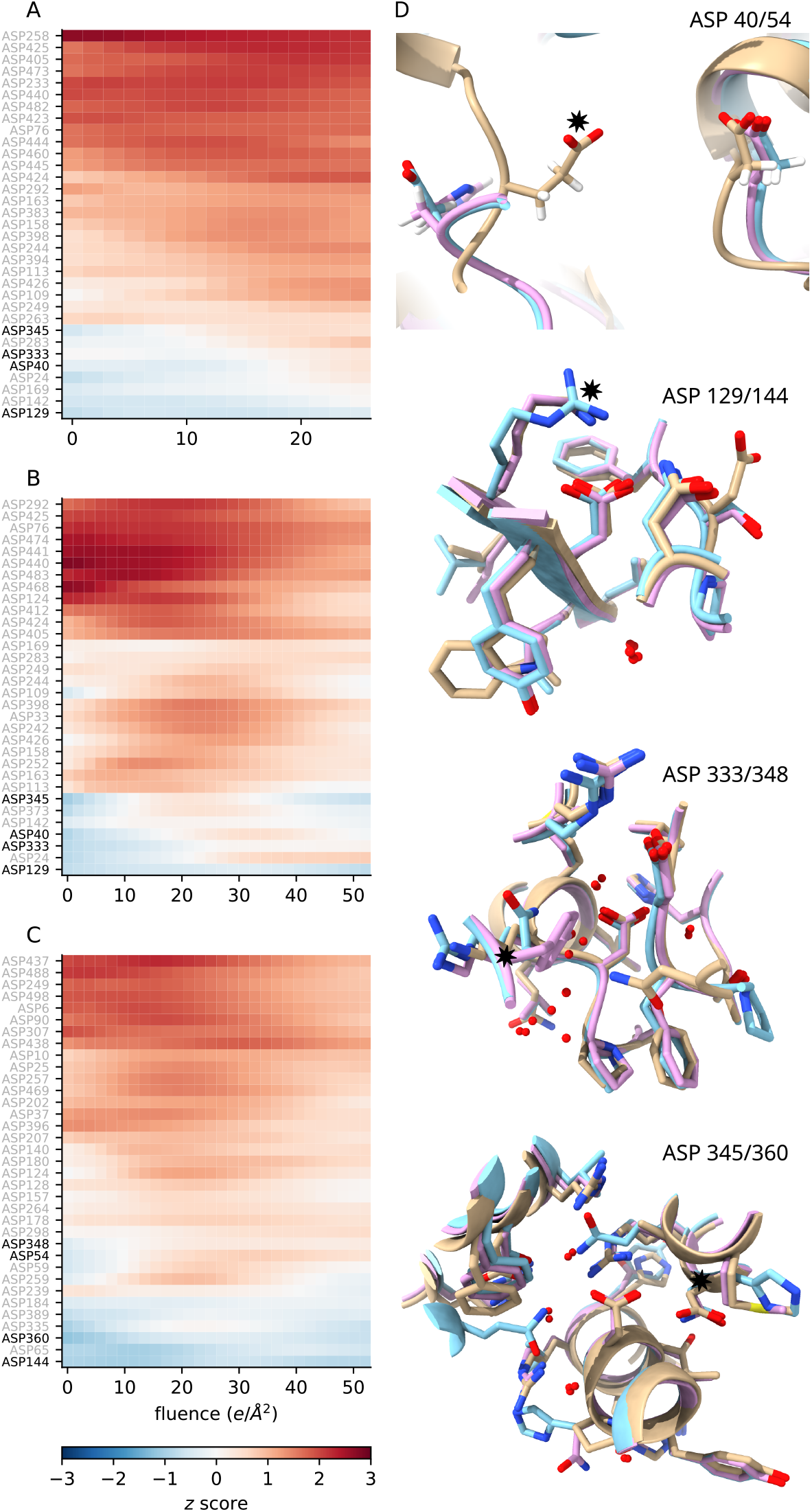
Environment dependence of RD to Asp carboxylates. (A–C) Heat maps showing behavior of *B*-factors of all Asp carboxylates in the three catalase structures as a function of fluence. Each square represents the *z*-score of C*γ* in the indicated Asp residue. (A) Catalase L, (B) catalase T, (C) catalase H. (D) Examples of Asp residues conserved across the three catalase structures. Catalase L is shown in cyan, catalase T in magenta and catalase H in beige. The residue given in the label is in the center of the image, other than in the top panel, where it is on the right. Stars indicate (top to bottom) catalase H Glu420, catalase L Arg120/catalase T Lys120/catalase H Lys135, catalase L Gln418/catalase T Tyr418, catalase L Asn51/catalase T Asn51/catalase H Asp65.

We chose four aspartates that are (a) conserved across the three catalases and (b) ordered at the start of imaging for further analysis. When reporting the results, we will use residue positions as given in the Protein Data Bank (PDB), giving the positions for catalase L/catalase T (which are the same) and catalase H separated by a slash. For example, Asp40/54 refers to Asp40 in catalase L and catalase T and to Asp54 in catalase H. We identified two residues (Asp40/54 and Asp 333/348) that are more susceptible to damage in catalase H than in catalase L and catalase T, one residue (Asp345/360) that is more susceptible in catalase L and catalase T than in catalase H, and one residue (Asp129/144) that is anomalously resistant to damage in all three structures. We compared the local environments of the four residues (Figure 3D). We used the package *PROPKA* [39], an empirical p*K* _a_ calculator, to quantify the effect of interactions of each residue with its local environment. The package also calculates the contribution of each interaction to the p*K* _a_, allowing us to assess its relative importance.

For all four residues considered, *PROPKA* identified key differences between the local environments in the three catalase structures. Starting with Asp40/54, *PROPKA* predicts an upwards shift in the p*K* _a_ of this functional group in catalase H compared to catalase L and catalase T, explained by an unfavorable Coulomb interaction with Glu420 in catalase H. On the other hand, in Asp333/348 the increased susceptibility in catalase H appears to be related to the presence of a solvent-accessible cavity next to the carboxylate, which is plugged by Gln418 and Tyr418 in catalase L and catalase T, respectively. The most notable feature of Asp345/360 indicated by *PROPKA* is the coupling between the protonation states of Asp345 and Asp65 in catalase H. The latter is predicted to be protonated (p*K* _a_ = 12.2), and is replaced by Asn51 in catalase L and catalase T. However, in this case the generally poor conservation of the local environment makes comparison difficult. Finally, the environment of Asp129/144 dominated by hydrogen bonding between the carboxylate and a positively charged side chain (Arg in catalase L, Lys in catalase T and catalase H). The identity of the positively charged amino acid does not appear to influence susceptibility to damage.

As a simple validation of our observations, we calculated the value of the raw cryo-EM maps at the coordinates of all Asp C*γ* atoms in the three catalase structures (Extended Data Figure 3). The maps were normalized before calculation to have zero mean and unit variance. The map values at the coordinates of Asp129/144 C*γ* were then compared to the average of the values for all aspartates. We found that the map value for Asp129/144 C*γ* is consistently higher than the average, suggesting it is indeed more resistant to damage than other Asp carboxylates.

### Modeling global RD using the Langevin equation

We have shown that our method can be used to analyze beam-induced changes in the sample on scales ranging from individual amino acid side chains to the whole macromolecule. Using this new information, we may try to model the non-linear behavior of *B*-factors described above. Several explanations for such behavior are possible, including large-scale rearrangement of the specimen or artifacts caused by the microscope hardware. However, here we explore whether the non-linearity may be explained by the dynamics of the protein itself.

Recall that a volume reconstructed by cryo-EM SPA represents the average of a population of particles. The quantity Δ*B*(*t*) is therefore proportional to the mean squared displacement (MSD) of each atom from its starting position. A given observed Δ*B*(*t*) may be rationalized by supposing that the time-dependent change in position of each atom in the population is governed by a stochastic process. The MSD of the stochastic process should have the same form as Δ*B*(*t*). The class of process we consider is Langevin dynamics, in which the motion of each atom in the macromolecule is assumed to be independent and is driven by the combination of a fluctuating force exerted on the atom by the surrounding medium and a drag force [17]. We further assume that atoms may be modeled as free particles, meaning there is no external potential. Langevin dynamics has the appealing property that in the overdamped regime (when the drag force is large) the MSD and hence Δ*B*(*t*) are a linear function of time, which replicates observations from XRD experiments [8]. On the other hand, in this study the *B*-factors increase non-linearly. We suggest that this non-linearity may be explained by Langevin dynamics in the underdamped regime. In this regime, the drag force is similar in magnitude to the random driving force, causing the process to retain memory of the atomic motions between successive stochastic impulses.

A derivation of the behavior of the quantity Δ*B*(*t*) predicted by Langevin dynamics is given in the Supplementary Information. The derived expression was fitted to the mean of Δ*B*(*t*) for each structure using non-linear least squares. Only data at fluences below 40 eÅ^-2^ were used for fitting. The reason for excluding high-fluence atomic models for the catalase H and catalase T datasets is that their *B*-factor distributions show non-monotonic behavior in this range. We attribute such anomalous behavior to difficulties in estimating *B*-factors for individual atoms after significant RD to the sample.

Values of Δ*B*(*t*) and fits of the function predicted by the Langevin equation are given in Figure 4, showing good agreement between the observations and the fitted function. Small deviations from the predicted values can be observed early in the exposure for the apoferritin and Dps time series, which may reflect the influence of intermittent charging artifacts at low fluence invoked by the authors of [15]. Nonetheless, the analysis overall suggests that the progressive disordering of atoms due to global RD may potentially be described by underdamped Langevin dynamics. The underdamping gives rise to a two-phase behavior, where the MSD scales as *t*^2^ at small fluence and *t* at large fluence. Although we concentrate on the mean Δ*B*(*t*), in practice Δ*B*(*t*) has a different value for each atom in the structure and thus is itself a random variable. The variation in Δ*B*(*t*) corresponds to variation in the MSD between atoms in the macromolecule and reflects differences in their local environment.

**Figure 4:**
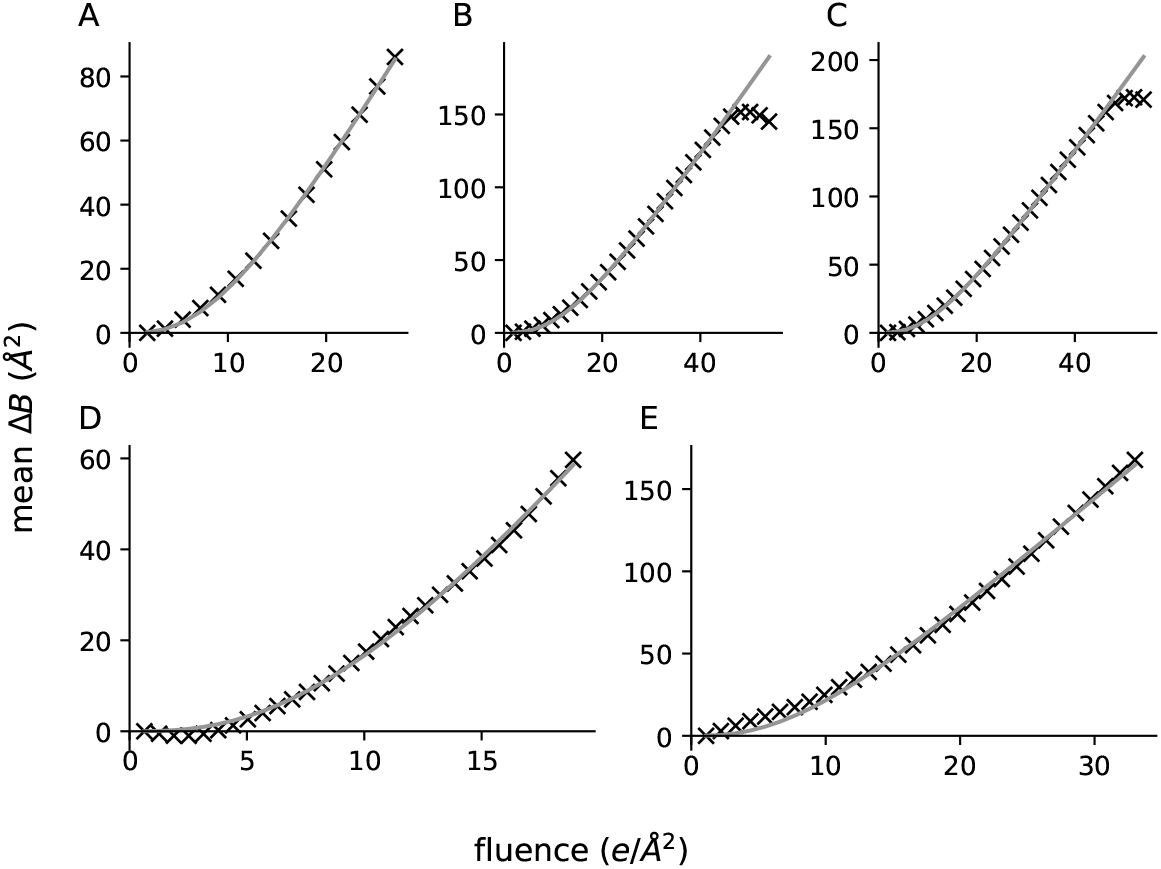
A stochastic process model of RD. In each panel, black crosses represent the mean difference of atomic *B*-factors from their values at the start of the time series, while gray lines are the values of this quantity predicted by the Langevin equation, with parameters of the equation fitted to the observations by non-linear least squares. (A) Catalase L, (B) catalase T, (C) catalase H, (D) apoferritin, (E) Dps.

## Discussion

In this contribution, we develop a time-resolved atomic model refinement algorithm which infers a ‘movie’ of protein structures from a time series of cryo-EM reconstructions at increasing electron fluence. Our algorithm takes into account correlations between the signal at different points in the time series, ensuring that the protein structure changes smoothly even at very high fluence, producing an atomic model that is as interpretable as one inferred from a standard dose-weighted cryo-EM reconstruction. We apply our algorithm to five cryo-EM datasets and identify a set of unifying trends. We show that the molecule expands and becomes increasingly disordered during imaging, and that the increase in disorder can be explained by a stochastic process acting on individual atoms. Exploiting the fact that our program can infer the time-dependent behavior of every atom in the molecule, we analyze the radiation sensitivity of specific amino acid side chains, showing that radiation-induced reactions observed in XRD at cryogenic temperatures are also relevant in cryo-EM SPA. Finally, we investigate the effect of perturbations in the local environment of Asp residues on their susceptibility to damage.

The beam-induced expansion that we observe has not previously been seen in cryo-EM and may have consequences for EM data processing. A similar expansion has been reported previously in two separate XRD studies [40, 54]. We can now confirm that this phenomenon also occurs in cryo-EM SPA, at least in the proteins which we examined in this study, which are highly symmetric and therefore have a well-defined center. A possible cause is radiolysis of the protein [22], which produces molecular fragments further away from one another than moieties joined by covalent bonds. Although the expansion effect is small, in high resolution cryo-EM datasets averaging of images over dose without taking into account the expansion (as is currently the case) may result in blurring of signal near the protein surface.

The primary effect of RD on protein structures is an increase in the disorder of atoms, which manifests itself as an increase in *B*-factors of the atomic model. We show that the increase in the average value of *B*- factors (Figure 1C/D) can be explained by an accumulation of individual damage events, and we show that these events can be modeled by Langevin dynamics. The motion appears to happen in an underdamped regime, giving rise to an accelerating damage rate at small time scales. The acceleration we observe is an example of the ‘latent dose’ effect, which has been reported numerous times in the EM literature as an apparent resistance of the sample to irradiation up to a certain dose [18]. The global aspect of RD fundamentally limits the dose that can be used in cryo-EM SPA experiments, and hence the achievable resolution. Therefore, we suggest that understanding the stochastic dynamics of RD in the sample is a fruitful avenue for future work which may give rise to methods to protect the macromolecule from RD. Furthermore, in proposing a physical model of RD we contribute to a broader theory of fluence-dependent information loss in cryo-EM SPA (referred to in [15] as the ‘grand scheme’). A pseudo-Brownian diffusion of water molecules is an important known component of this theory [34], and could be integrated into our stochastic process model in future.

Although global effects are key to understanding the nature of the damage process, it is site-specific damage that is of most concern when drawing biological conclusions from cryo-EM reconstructions. Regarding the atomic coordinates, we find that the difference from the dose-weighted structure is very small, with a structure RMSD well below 1 Å. This is good news for experimental structure determination, as it suggests that site-specific damage does not significantly affect map interpretation at resolutions currently attainable by cryo-EM. We do not exclude, however, that certain moieties may be so sensitive to exposure to the electron beam that further care is warranted. Indeed, one such case (a bacterial photosystem II) has been described in the literature [28]. We suggest that when there is doubt about whether a feature in a cryo-EM map is caused by damage, application of our atomic model refinement scheme could clarify the origin of the observation. Regarding *B*-factors, this study confirms that patterns of site-specific damage documented by XRD are broadly applicable in cryo-EM SPA. In particular, we note that carboxylate groups on Glu and Asp that are initially ordered become disordered over time at a rate much faster than comparable atoms in other functional groups. So far, explanations of the poor visibility of Asp/Glu side chains in cryo-EM reconstructions have focused on charge effects [33, 53, 59], however, our analysis indicates that radiation-induced decarboxylation must also be taken into account when interpreting EM data.

We find that certain Asp side chains show anomalous resistance to radiation, and, exploiting the fact that the three catalase structures examined in this work have a high degree of sequence similarity, we attempt to relate changes in the local environment of Asp to its damage susceptibility. The reaction scheme for radiation-induced decarboxylation conveniently summarized in [12] provides a way to rationalize the effects we observe. The radiolysis reaction proceeds by the stabilization of an electron hole on the carboxylate group, creating an unstable one-electron oxidized intermediate which is immediately decarboxylated. There is also a side reaction involving the formation of a one-electron reduced Asp^•–^ radical that is then protonated [1]. The side reaction does not lead to decarboxylation. In the environments shown in Figures 3A/B, we see that an adjacent positive charge has a stabilizing effect on the carboxylate, whereas an adjacent negative charge has a destabilizing effect. The former agrees with a previous study of RD susceptibility in XRD [19]. We speculate that surrounding positive charge may prevent hole stabilization on the carboxylate moiety. The behavior of Asp345/360 is more difficult to rationalize, however we suggest that the coupling of Asp360 and Asp65 in catalase H has a protective effect by allowing rapid proton transfer to Asp360, thus favoring the reductive side reaction mentioned previously. Finally, in the environment of Figure 3C, exposure of the side chain to solvent appears to increase damage susceptibility. Similar effects have previously been explained by a reduction in the steric hindrance of the carbon dioxide molecule produced during radiolysis [11]. CO_2_ must adopt a planar conformation to fully dissociate from the protein, and surrounding bulky side chains may prevent this from happening. Quantum-chemical calculations carried out in different atomic environments may help validate the mechanisms we mention.

Finally, we point out that the atomic model refinement algorithm described in this work is not fundamentally limited to RD time series. By changing the covariance model (and possibly rebuilding the atomic model to best describe the observations at different time points), it would for example be possible to refine structures into series of reconstructions obtained by time-resolved cryo-EM [4], or imaged under different conditions. Our algorithm represents a parallelizable paradigm for approaching these and similar optimization problems.

## Data availability

The source code of an implementation of the atomic model refinement algorithm is available on GitHub (https://github.com/as2875/sffit). Atomic model time series are available on Zenodo (https://zenodo.org/records/21107538).

## Acknowledgments

We thank R. Henderson for his comments on the manuscript, and we thank C.J. Russo and S.H.W. Scheres for helpful discussions and advice. We also thank J. Grimmett, T. Darling, and I. Clayson for access to scientific computing resources. This work was supported by the Medical Research Council under Grant No. MC UP A025 1012 to G.N.M. H.W. is supported by the Astex Pharmaceuticals Sustaining Innovation postdoctoral scheme.

## Author contributions

A.S. designed the algorithm, implemented the algorithm, analyzed the results, and formulated the damage model. H.W. produced the catalase reconstructions and formulated the damage model. G.N.M. conceived the study and supervised the study. A.S. wrote the manuscript with input from all authors.

## Methods

### Probabilistic model

We produce a tractable algorithm by formulating multiple-model refinement as a variational inference (VI) problem, based on the idea of approximating the true data-generating process by one that is parametrized by a series of atomic models. The algorithm therefore has two components: a model of the data-generating process, and a procedure for fitting the variational approximation.

The data-generating process in question should describe the ESP of a macromolecule (averaged over the ensemble of particles used for the reconstruction) as a function of fluence. Instead of modeling the ESP directly, we model its Fourier transform, as is common in cryo-EM data processing. Fourier-space models are convenient because the observation noise in different Fourier coefficients may be assumed independent [35], moreover, the ESP and its Fourier transform are easily interconvertible since the latter is a linear operator. We model the time-dependent behavior of each Fourier coefficient in the reconstruction as a Gaussian process (GP), a stochastic process in which any finite collection of values at fixed time points follows a multivariate normal distribution (see [42] for a comprehensive introduction to GPs). We further suppose that observations of the Fourier coefficients are corrupted by additive independent and identically distributed (iid) Gaussian noise.

Formally, suppose there are *K* cryo-EM maps in the time series. We represent by **z** ∈ ℂ^*K*^ the vector containing the ‘true’ values of a single Fourier coefficient at each time point and by **x** ∈ ℂ^*K*^ an analogous vector containing the observed values of each Fourier coefficient, which are the true values corrupted by additive noise. We place a GP prior on **z**. For the purposes of specifying resolution-dependent hyperparameters, the Fourier coefficients are divided into resolution bins, a heuristic which is common in structure determination by cryo-EM and XRD. The prior and likelihood are then

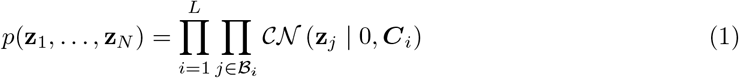

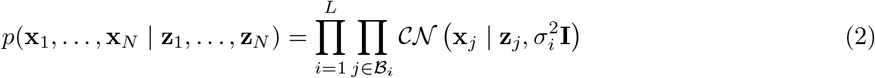

where **C**_*i*_ is the *K*×*K* covariance matrix of the prior, **I** is the identity and 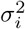 is the variance of the observation noise. *L* is the number of resolution bins and the notation *j* ∈ ℬ_*i*_ denotes membership of a Fourier coefficients in resolution bin *i. CN* denotes a complex normal distribution with iid real and imaginary parts. Note that the likelihood assumes that the observation noise is constant over time, which is a valid assumption as long as the fluence per frame is constant. A posterior distribution derived from the prior and likelihood is required to make Bayesian inferences about **z**_*i*_. For GPs, the posterior has a closed form,

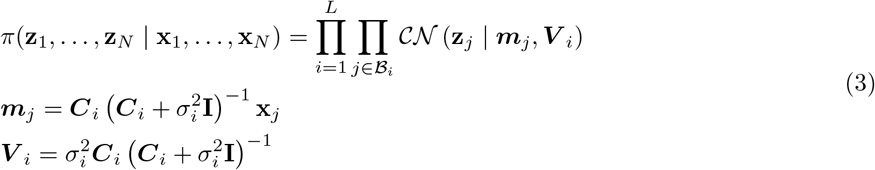

### Variational approximation

Our aim is to approximate the true posterior in Equation 3 by a distribution with a mean given by Fourier coefficients **y**_*j*_(Θ) calculated from an atomic model with parameters Θ,

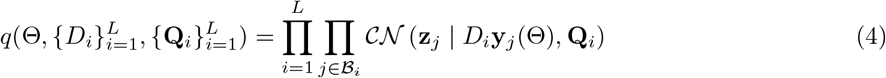

where *D*_*i*_ is a constant that scales the atomic model onto the observations and **Q**_*i*_ is the covariance of the variational distribution. The parameters of the approximation Θ, *D*_*i*_ and **Q**_*i*_ are found by minimizing the Kullback-Leibler (KL) divergence *D* [*π*|| *q*] := *D* (Θ, {*D*_*i*_}, {**Q**_*i*_}). Note we use the forward KL divergence instead of the reverse KL generally encountered in VI [6]. This choice makes our objective more similar to the one used by the popular atomic model refinements package *REFMAC* [36] and *Servalcat* [58]. For our probabilistic model, the KL divergence has the form

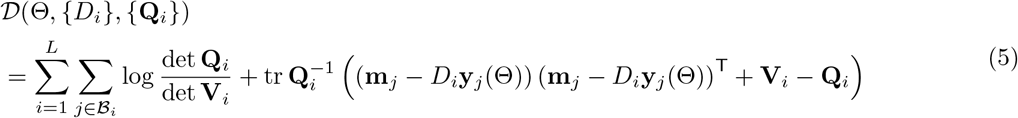

where **m**_*i*_ and **V**_*i*_ may be calculated using Equation 3 and ^T^ is the conjugate transpose. In practice, we restrict **Q**_*i*_ to the manifold of diagonal-plus-rank-one matrices as this approximation captures the shape of the underlying distribution while having the advantage that simple formulas exist for the inversion of such matrices.

### Estimation of distributional parameters

We will now describe a procedure for estimating *D*_*i*_, **Q**_*i*_ and Θ given posterior means and variances calculated using Equation 3. We return to the problem of estimating **C**_*i*_ and 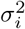 later. The objective in Equation 5 is similar to the objective function used by the software package *REFMAC*, which reduces to a non-linear least squares problem in Θ if the linear parameters *D*_*i*_ and **Q**_*i*_ are known. Therefore, similarly to *REFMAC*, we apply block coordinate descent to minimize the function with respect to *D*. The algorithm alternates between (1) optimization with respect to the linear parameters while Θ is kept fixed and (2) optimization with respect to Θ while the linear parameters are kept fixed. It may be shown that the optimal values for *D*_*i*_ and **Q**_*i*_ for a given Θ satisfy

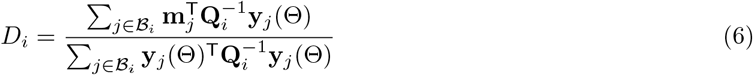

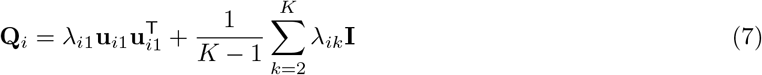

where *λ*_*ik*_ and **u**_*ik*_ are the *k*th eigenvalue and corresponding eigenvector (in descending order) of the matrix

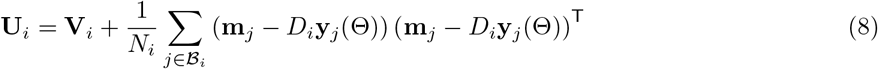

Derivations of the parameter updates are given in the Supplementary Information. Here, *N*_*i*_ is the number of Fourier coefficients in resolution bin *i*. Equation 8 may be interpreted as combining two sources of uncertainty: the first term is the covariance of the posterior expectation **m**_*i*_ under the GP, while the second term is the covariance of the model residuals, representing signal in the map unaccounted for by the atomic model. At each iteration, the algorithm finds the optimal values for *D*_*i*_ and **Q**_*i*_ by alternating between the updates in Equations 6 and 7 (with Θ fixed) until convergence.

### Optimization scheme

As noted above, optimization with respect to Θ is a generalized non-linear least squares problem

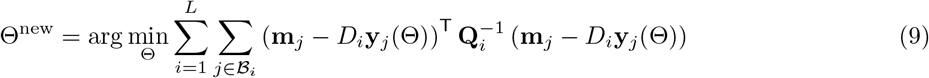

Instead of directly solving Equation 9, we perform one majorization-minimization (MM) update at each iteration. MM is an iterative optimization scheme which has been extensively reviewed elsewhere [50]. Briefly, an MM parameter update consists of (1) constructing a surrogate function which majorizes the objective at the current estimate of the parameters and (2) optimizing the surrogate with respect to the parameters. A function *f* (*x*) is said to majorize *g*(*x*) at *y* when *f* (*x*) ≥ *g*(*x*) for all *x* and *f* (*y*) = *g*(*y*). The surrogate function may be chosen to have desirable properties, such as being separable in the parameters.

It may be shown that Equation 10 below gives a valid MM update (derivations are given in the Supplementary Information).

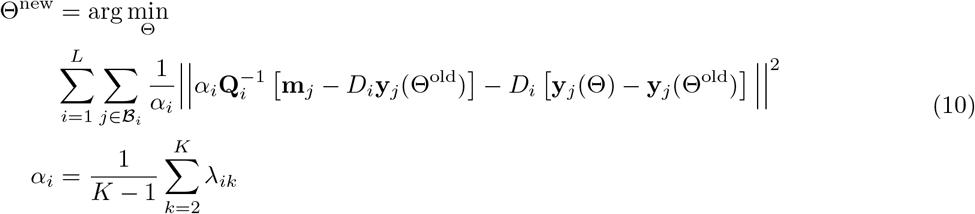

|| · || denotes the *L*_2_ norm. Note that this objective function is block-separable so optimization can be performed independently with respect to the parameters of each atomic model in the time series, a task which we perform using the program *Servalcat*. In practice, as in standard model refinement, restraints on bond lengths, angles and *B*-factors must be used to obtain chemically reasonable atomic models; the restraints are described in the Supplementary Information.

### Prior covariance

We now return to the problem of specifying a prior covariance. A goal of this work is to use the time series of atomic models produced by the program to formulate a stochastic process model describing RD in cryo-EM. The prior should therefore not assume a time dependence of the atomic model parameters. In this work, we use a scaled radial basis function (RBF) covariance. The RBF covariance is a popular GP covariance function and imposes infinite differentiability on the underlying stochastic process [56].

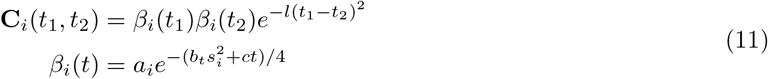

*l* controls the correlation length, *b*_*t*_ is a scaling *B*-factor that is different for each time point and *s*_*i*_ is the resolution in Å^-1^ of bin *i*. The term *ct* in the exponent may be interpreted as modeling mass loss from the sample over time. The scaling term *β*(*t*) is identical to that used by *RELION* to calculate radiation dose-weighted reconstructions [60], with the difference that we impose a linear time dependence on the mass loss term instead of fitting a separate parameter for each time point. We find this is required to avoid overfitting. The parameters of the covariance, as well as the noise variance 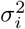, are estimated by a combination of maximum likelihood and spectral estimators (details are given in the Supplementary Information).

### Datasets

We applied our algorithm to five cryo-EM reconstruction time series. The datasets were processed to obtain a series of reconstructions of the macromolecular ESP, with each reconstruction using data from a single frame of all movies in the dataset [37].

Three of the five datasets are reconstructions of catalase enzymes from different species: *Rhizobium radiobacter* (which we will refer to as catalase T), *Micrococcus luteus* (catalase L) and human erythrocyte catalase (catalase H) [47]. Dose-weighted reconstructions calculated from the catalase datasets have a nominal resolution in the range 1.7–1.9 Å. The two bacterial catalases have a sequence identity of 69%, while catalase H has 47% and 44% sequence identity with catalase T and catalase L, respectively. Catalase is a heme-containing protein that catalyzes the dismutation of hydrogen peroxide into oxygen and water, thus forming part of the cellular defense against oxidative damage [13]. The fourth dataset contains reconstructions of *Mus musculus* heavy chain apoferritin. The dose-weighted reconstruction calculated from the dataset has a nominal resolution of 2.2 Å. The fifth time series consists of reconstructions of *E. coli* Dps, a ferritin homolog with iron binding and ferroxidase activity [25]. The dose-weighted reconstruction calculated from the Dps dataset has a nominal resolution of 2.4 Å. The total fluence of micrographs in the Dps dataset was 55 eÅ^-2^, however, only data up to a fluence of 33 eÅ^-2^ were used in the analysis below, as reconstructions calculated at higher fluence were found to have SNR too low to allow atomic modeling. The apoferritin and Dps datasets were originally published in [15].

All datasets we used were collected on HexAuFoil grids, which reduce beam-induced particle movement during imaging to less than 1 Å [37]. The catalase data were collected on HexAuFoil grids with 300 nm holes, while the hole diameter was 200 nm and 100 nm for the apoferritin and Dps datasets, respectively.

---

**Extended Data Figure 1:**
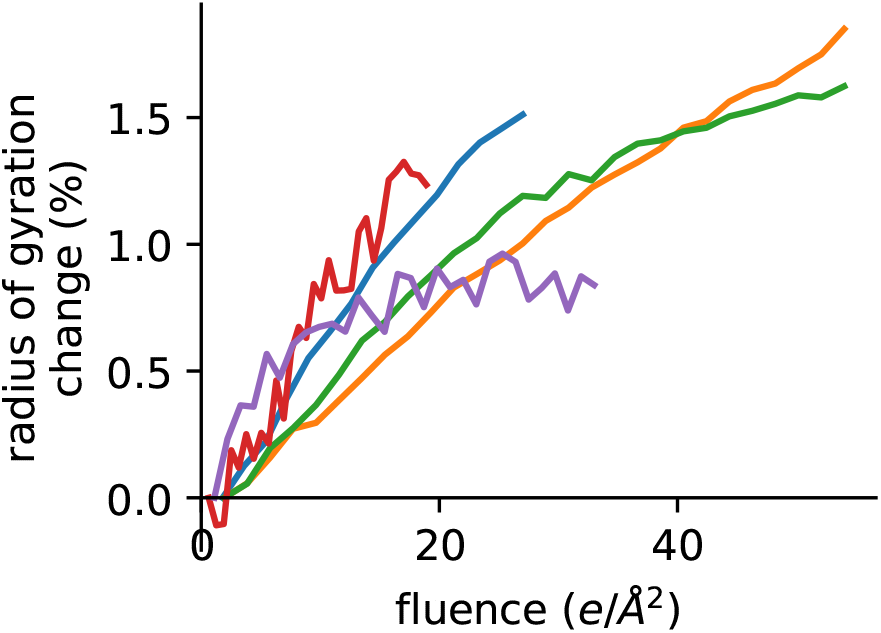
Radius of gyration calculated from time series of cryo-EM reconstructions. *y* -axis shows percentage difference between radius of gyration at indicated fluence and radius of gyration for the first map in the time series. Maps were scaled to have the same power spectrum before calculation of the radius of gyration. Colored lines indicate different datasets, colors are as in Figure 1.

**Extended Data Figure 2:**
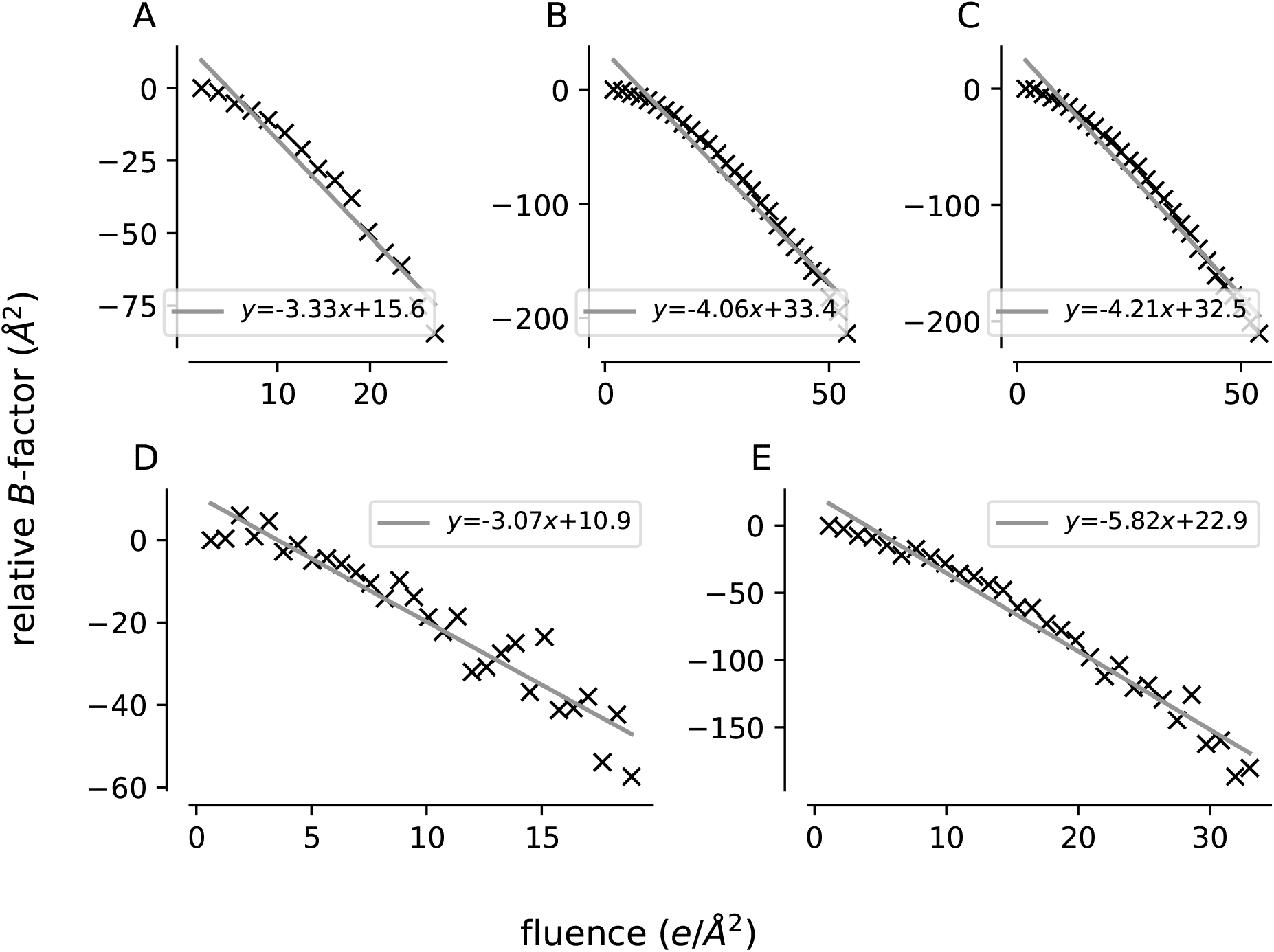
Relative *B*-factors calculated from time series of cryo-EM reconstructions using the formulas derived in [45]. *B*-factors are relative to the first reconstruction in the time series. Data are shown as black crosses, gray lines are linear functions fitted by ordinary least squares. (A) Catalase L, (B) catalase T, (C) catalase H, (D) apoferritin, (E) Dps.

**Extended Data Figure 3:**
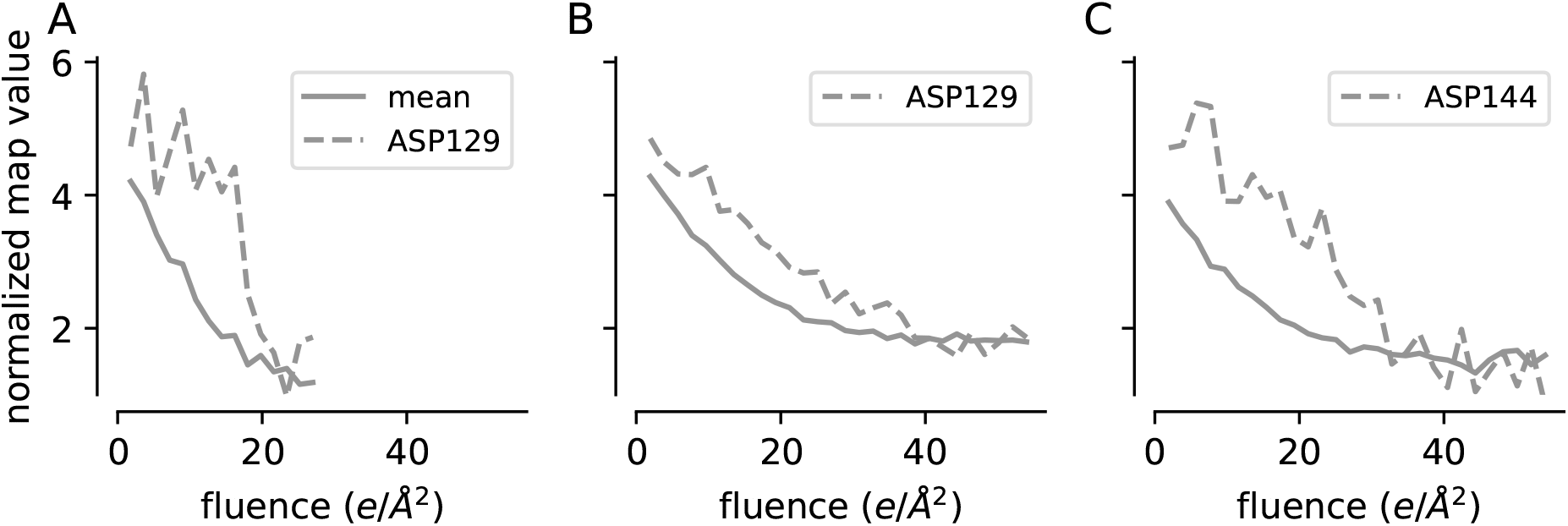
Normalized values of catalase maps at the locations of Asp C*γ* atoms as a function of fluence. Maps were normalized to have zero mean and unit variance before calculation. In each panel, the solid line represents the mean of this value evaluated for all Asp residues in the structure and the dashed line represents the value at the location of the C*γ* atom of the indicated residue. (A) Catalase L, (B) catalase T, (C) catalase H.

