## Supplementary Information for "Atomic modeling of radiation damage in cryoelectron microscopy datasets"

#### 1 Derivation of stochastic process model

If the motion of a particle with mass $m$ is governed by Langevin dynamics, the infinitesimal changes in its position $\mathbf{r}$ and momentum $\mathbf{p}$ are given by

$$\mathrm{d}\mathbf{r} = \frac{\mathbf{p}}{m}\mathrm{d}t \tag{S1}$$

$$\mathrm{d}\mathbf{p} = \left(-\nabla U(\mathbf{r}) - \gamma\mathbf{p}\right)\mathrm{d}t + \sqrt{2m\gamma k_B T}\mathrm{d}\mathbf{W} \tag{S2}$$

where $U(\mathbf{r})$ is a potential, $\gamma$ is the coefficient of friction, $k_B$ is the Boltzmann constant, $T$ is temperature and $\mathbf{W}$ is Brownian motion. In what follows, we assume that there is no external potential, i.e. $U(\mathbf{r}) = 0$. In a molecular model, the external potential represents the influence of surrounding atoms, for example through covalent bonds, on the movement of the atom under consideration. Electron energy loss spectra of proteins have their largest peak around 20 eV, in the plasmon loss region [7]. On the other hand, the dissociation energy of a typical organic covalent bond is 3–4 eV [2], and structural rearrangements may take place at smaller energies. The magnitude of the energy imparted by the electron relative to the dissociation energy suggests that the latter may be ignored as a first approximation.

Given the assumptions in the previous paragraph, it may be shown that under the initial condition $\mathbf{r} = 0, \mathbf{p} = 0$ at $t = 0$ the probability density in phase space is given by [9]

$$\mathbf{r}, \mathbf{p} \mid t \sim \mathcal{N}\left(0, \begin{bmatrix} \sigma_r^2 & \sigma_{rp} \\ \sigma_{rp} & \sigma_p^2 \end{bmatrix} \otimes \mathbf{I}_3\right) \tag{S3}$$

where $\otimes$ is the Kronecker product, $\mathbf{I}_3$ is the $3 \times 3$ identity matrix and

$$\sigma_r^2 = \frac{k_B T}{m\gamma^2}\left(1 + 2\gamma t - \left(2 - e^{-\gamma t}\right)^2\right) \tag{S4}$$

$$\sigma_p^2 = m k_B T\left(1 - e^{-2\gamma t}\right) \tag{S5}$$

$$\sigma_{rp} = \frac{k_B T}{\gamma}\left(1 - e^{-\gamma t}\right)^2 \tag{S6}$$

We are interested in the marginal distribution of $\mathbf{r}$ at time $t$. Since the joint distribution is Gaussian, the marginal distribution is also Gaussian and is given simply by the corresponding block of the covariance matrix of the joint distribution [1].

$$\int_{\mathbf{p}\in\mathbb{R}^3} p(\mathbf{r}, \mathbf{p}, t)\mathrm{d}\mathbf{p} = \mathcal{N}\left(0, \sigma_r^2\mathbf{I}_3\right) \tag{S7}$$

The variance of the marginal distribution is proportional to the quantity $\Delta B(t)$ described in the main article. Non-linear least squares was used to fit the parameters $\gamma$ and $k_B T/m$ to observed values of $\Delta B(t)$ for each of the five structures analyzed in the main article.

The linear model of $B$-factor dependence on time which is commonly used in XRD [3] may be recovered from Langevin dynamics by considering the overdamped regime, that is for large $\gamma$. In this case, $\sigma_r^2 \approx \frac{2k_B T}{m\gamma}t$, meaning the variance of position increases linearly with time, as required.

#### 2 Derivation of model parameter updates

In this section, we derive the block coordinate descent parameter updates for $D_i$, $\mathbf{Q}_i$ and $\Theta$. To derive the update for $D_i$, differentiate the objective in Equation 5 with respect to this parameter:

$$\frac{\partial \mathcal{D}}{\partial D_i} = 2 \sum_{j \in \mathcal{B}_i} D_i \mathbf{y}_j^\mathsf{T} \mathbf{Q}^{-1} \mathbf{y}_j - \mathbf{y}_j^\mathsf{T} \mathbf{Q}^{-1} \mathbf{m}_j \tag{S8}$$

Equation 6 is obtained by finding the root of the expression for the derivative.

To derive the update for $\mathbf{Q}_i$, write the objective in the form

$$\mathcal{D}\left(\Theta_i, \{D_i\}, \{\mathbf{Q}_i\}\right) = N_i \sum_{i=1}^{L} \log\det \mathbf{Q}_i + \operatorname{tr} \mathbf{Q}_i^{-1} \mathbf{U}_i + \text{const.} \tag{S9}$$

where $\mathbf{U}_i = \mathbf{V}_i + \frac{1}{N_i} \sum_{j \in i} \left(\mathbf{m}_j - D_i \mathbf{y}_j(\Theta)\right) \left(\mathbf{m}_j - D_i \mathbf{y}_j(\Theta)\right)^\mathsf{T}$ and $N_i$ is the number of coefficients in the $i$th resolution bin. Suppose $\mathbf{Q}_i$ has the eigenvectors $\mathbf{u}_{i1}, \mathbf{u}_{i2}, \ldots, \mathbf{u}_{iK}$ and eigenvalues $\lambda_{i1}+d, \lambda_{i2}+d, \ldots, \lambda_{iK}+d$. We require $\mathbf{Q}_i$ to be a positive definite diagonal-plus-rank-$n$ matrix, meaning $\lambda_{ik} > 0$ for $k = 1 \ldots n$ and $\lambda_{ik} = 0$ for $k = n+1 \ldots K$. The objective in Equation S9 is identical up to scaling to the objective used in probabilistic PCA [12]. In [12], it is shown that the optimum value of the objective is obtained when $\lambda_{i1} \ldots \lambda_{in}$ are the $n$ largest eigenvalues of $\mathbf{U}_i$, $\mathbf{u}_{i1} \ldots \mathbf{u}_{in}$ are the corresponding eigenvectors and $d = \frac{1}{K-n} \sum_{k=n+1}^{K} \lambda_{ik}$. We use $n = 1$ in the main article.

The MM update for $\Theta$ makes use of the following result: the quadratic form $f(\mathbf{v}) = \mathbf{v}^\mathsf{T} \mathbf{A} \mathbf{v}$ is majorized at $\mathbf{v}'$ by [11]

$$\begin{aligned} g(\mathbf{v}, \mathbf{v}') &= \mathbf{v}^\mathsf{T} \mathbf{M} \mathbf{v} - 2\mathbf{v}^\mathsf{T} (\mathbf{M} - \mathbf{A}) \mathbf{v}' + \mathbf{v}' T (\mathbf{M} - \mathbf{A}) \mathbf{v}' \\ &= \left(\mathbf{v} - \left(\mathbf{I} - \mathbf{M}^{-1}\mathbf{A}\right) \mathbf{v}'\right)^\mathsf{T} \mathbf{M} \left(\mathbf{v} - \left(\mathbf{I} - \mathbf{M}^{-1}\mathbf{A}\right) \mathbf{v}'\right) + \text{const.} \end{aligned} \tag{S10}$$

if $\mathbf{M} \succeq \mathbf{A}$. Terms independent of $\mathbf{v}$ are omitted in the second line. Equation 10 is obtained by setting $\mathbf{A} = \mathbf{Q}_i^{-1}$, $\mathbf{v} = \mathbf{m}_j - D_i \mathbf{y}_j(\Theta)$, $\mathbf{v}' = \mathbf{m}_j - D_i \mathbf{y}_j(\Theta^{\text{old}})$ and $\mathbf{M} = \frac{1}{K-1} \sum_{k=2}^{K} \lambda_{ik} \mathbf{I}$.

#### 3 Estimation of covariance hyperparameters

The model covariance $\mathbf{C}$ is estimated from the data by maximizing the marginal likelihood function. The marginal likelihood is the probability of the observations given the hyperparameters, obtained by integrating over values of the GP prior [8]. The marginal log-likelihood is given by

$$\begin{aligned} \log p\left(\mathbf{x}_1, \ldots, \mathbf{x}_N \mid \left\{\mathbf{C}_i, \sigma_i^2\right\}_{i=1}^{L}\right) &= \log \prod_{i=1}^{L} \prod_{j \in \mathcal{B}_i} \mathcal{CN}\left(\mathbf{x}_j \mid 0, \mathbf{C}_i + \sigma_i^2 \mathbf{I}\right) \\ &= \sum_{i=1}^{L} \sum_{j \in \mathcal{B}_i} -K \log \pi - \log\det\left(\mathbf{C}_i + \sigma_i^2 \mathbf{I}\right) - \mathbf{x}_j^\mathsf{T} \left(\mathbf{C}_i + \sigma_i^2 \mathbf{I}\right)^{-1} \mathbf{x}_j \end{aligned} \tag{S11}$$

where $\mathcal{B}_i$ is the set of indices of Fourier coefficient in resolution bin $i$ and there are $L$ bins in total. We can rewrite the last term to be a function of the empirical covariance in each resolution bin,

$$\sum_{j \in \mathcal{B}_i} \mathbf{x}_j^\mathsf{T} \left(\mathbf{C}_i + \sigma_i^2 \mathbf{I}\right)^{-1} \mathbf{x}_j = \operatorname{tr}\left[\left(\mathbf{C}_i + \sigma_i^2 \mathbf{I}\right)^{-1} \sum_{j \in \mathcal{B}_i} \mathbf{x}_j \mathbf{x}_j^\mathsf{T}\right] \tag{S12}$$

The use of the empirical covariance reduces the memory requirements of fitting. We perform hyperparameter estimation by first binning Fourier coefficients by resolution, then calculating the empirical covariance in each resolution shell and finally finding the hyperparameters that maximize Equation S11. Smoothness of the parameters $b_t$ with respect to $t$ is ensured by L2-regularizing the second derivative of this quantity. Optimization is performed using L-BFGS [4]. In order to ensure the SNR of the maps is estimated correctly, we use spectral estimators of the signal and noise power ($a_i$ and $\sigma_i^2$ in our notation). Denoting by $\mu_{ik}$ the $k$th eigenvalue of the empirical covariance matrix of the $i$th resolution bin, the estimators are $\sigma_i^2 = \min_k \mu_{ik}$ and $a_i = \frac{-\sigma_i^2 + \sum_k \mu_{ik}}{\operatorname{tr} \mathbf{C}_i(a_i=1)}$. Here the denominator is the trace of the model covariance evaluated with unit power and other parameters equal to the optimum parameters determined by the maximum likelihood procedure outlined above.

### 4 Details of atomic model refinement

The surrogate objective in Equation 5 was optimized using the program *Servalcat* [14], which is designed to refine the parameters of atomic models of macromolecules against ESP maps. Site occupancies, which represent the proportion of molecules in the ensemble in which the atom is located at the corresponding atomic coordinate, are not refined in this work as all-atom occupancy refinement is not standard practice in experimental structural biology [13]. Refinement was performed using a $B$-factor restraint weight of 2.0 for the catalase and apoferritin structures and 1.0 for the Dps structure. Jelly-body and non-crystallographic symmetry restraints were also used. 20 MM updates were performed for the catalase and Dps datasets, 12 for the apoferritin dataset. In all cases, *Servalcat* was run for 200 cycles in the first update and 40 cycles in subsequent updates.

Starting atomic models for the catalase datasets were taken from [10]. PDB depositions 7A4M [5] and 6ZGL [6] were used as starting models for the apoferritin and Dps datasets, respectively. Atomic models of apoferritin and Dps were first rebuilt into the deposited dose-weighted maps (respectively Electron Microscopy Data Bank depositions EMD-51784 and EMD-51778).
